# Fetal protection against bovine viral diarrhea virus types 1 and 2 after vaccination of the dam with the DIVENCE vaccine

**DOI:** 10.1101/2024.04.12.589196

**Authors:** Ester Taberner, Marta Gibert, Carlos Montbrau, Irene Muñoz, Joaquim Mallorquí, Héctor Santo Tomas, Antoni Prenafeta, Ricard March

## Abstract

The aim of this study was to demonstrate the efficacy of DIVENCE, a vaccine against BVDV types 1 and 2 (BVDV-1 and BVDV-2) transplacental infection, following a booster regimen in heifers. Calves of two-to-three months of age were given two intramuscular doses three weeks apart and a booster vaccine six months later. Efficacy was evaluated by means of a challenge with virulent BVDV-1 or BVDV-2 administered via the intranasal route at 85 days of gestation. Clinical signs, serology, viral shedding, WBC count and viremia were monitored after the challenge. Sixty-six days post-challenge, the fetuses were assessed for BVDV to detect transplacental infection. The results demonstrate a reduction in hyperthermia, leukopenia, viral shedding, and viremia in vaccinated animals post-challenge with BVDV-1 and BVDV-2. Most importantly, DIVENCE administered prior to breeding protected 94% of the fetuses against BVDV transplacental infection overall across both challenge trials (BVDV-1 and BVDV-2).

## INTRODUCTION

Bovine viral diarrhea virus (BVDV) is one of the most important viral pathogens in cattle with a significant economic impact for the cattle industry (1,2,3,4). This bovine pestivirus is found worldwide; the proportion of BVDV-exposed herds ranges from 46% in Europe to 78% in Oceania (5). Only a few European countries have eradicated the virus (3,6). BVDV infection has a broad spectrum of manifestations, ranging from no clinical signs, to severe disorders of different organ systems (respiratory, digestive, reproductive, etc.), and even to death. Immunosuppression induced by acute BVDV infection predisposes animals to secondary or co-infections with other pathogens (7,8,9). The interaction between BVDV and secondary pathogens thus contributes to the development of the bovine respiratory disease (BRD) complex (10,11,12,13).

BVDV is a positive-strand RNA pestivirus with two antigenically distinct genotypes (types 1 and 2). Both genotypes have non-cytopathic (ncp) and cytopathic (cp) forms (biotypes), classified according to whether they produce visible changes in cell cultures (14). However, only ncp biotypes result in persistent infection; cp strains emerge by spontaneous mutation in animals with persistent ncp BVDV infection, leading to mucosal disease (15). A further classification into subgenotypes is needed due to genetic diversity. BVDV-1 can be classified into at least 22 subgenotypes (1a to 1u), and BVDV-2 into 3–4 subgenotypes (2a to 2c, 2d potentially). The most prevalent subgenotypes are 1a, 1b and 2a (16).

The virus can infect the female genital tract and cross the placenta, causing intrauterine infection of the fetus. If ncp BVDV infection occurs between Days 42-125 of gestation, the fetus will be born persistently infected (PI) (17). Approximately half of PI animals appear clinically normal; consequently, infection can be identified only through laboratory analyses (3). PI calves exhibit viral shedding, usually in large amounts over their entire life, and are considered the primary source for the spread of BVDV (18,19). Other reproductive outcomes in susceptible pregnant heifers and cows (transient infertility, embryonic death, abortions, stillbirth, congenital defects, and malformations) depend on the gestational stage of pregnancy (20).

Vaccination against BVDV is an effective tool for disease control, as it reduces the reinfection risk in the herd (21) as well as the likelihood of fetal infection, thereby also reducing the number of PI calves born in a herd (8). Subunit vaccines provide an opportunity to develop safer vaccines that allow infected and vaccinated animals to be differentiated (DIVA or marker vaccines). DIVA vaccines have been used in the veterinary field for decades. They carry at least one antigenic protein less than the wild-type virus; the diagnostic test thus measures the antibodies against the absent protein(s) to identify infected animals (22). A novel BVDV subunit vaccine has been developed (DIVENCE), comprising five different antigens – live-genetically modified BoHV-1 (double-deleted glycoprotein E and thymidine kinase, gE-/tk-), live-attenuated BRSV, inactivated PI3 virus, BVDV-1 E2 recombinant glycoprotein, and BVDV-2 E2 recombinant glycoprotein – designed to protect cattle against all these pathogens. This subunit vaccine aims to improve the efficacy and safety of current BVDV vaccines and also contributing as a DIVA vaccine.

The purpose of the study was to determine the efficacy of DIVENCE for fetal protection in vaccinated animals challenged with ncp BVDV-1 and BVDV-2 strains. In addition, changes in clinical signs, WBC count, viral shedding, and viremia were also assessed post-challenge.

## MATERIALS AND METHODS

### Animals and vaccination

All procedures involving cattle were approved by the local government (Generalitat de Catalunya, ref. 10352) and followed the recommendations of Directive 2010/63/EU of the European Parliament and Hipra’s Animal Health Ethical Review Board. Multiple-source heifers (n=59) were initially selected in the study. Heifers were two-to-three months of age and had no antibodies against BVDV (HerdCheck BVDV antibody test, IDEXX); in addition, they were confirmed to be free of persistent BVDV infection by real-time reverse transcription polymerase chain reaction (RT-qPCR). None of the heifers had serum neutralizing (SN) antibodies against BVDV-1 and BVDV-2 prior to the start of the study. Animals were randomly distributed into two challenge trials (BVDV-1 and BVDV-2).

For the BVDV-1 challenge, 29 heifers were randomly assigned to vaccination and control groups; 19 received two intramuscular (IM) doses of DIVENCE (2 mL) on Days 0 and 21 of the study followed by a booster vaccine six months later; the remaining ten heifer calves received sterile PBS following the same regimen.

For the BVDV-2 challenge, 30 heifers were randomly assigned to vaccination and control groups: 20 received two IM doses of DIVENCE (2 mL) on Days 0 and 21 of the study followed by a booster vaccine six months later; the remaining ten heifer calves received sterile PBS following the same regimen.

The vaccine was reconstituted and diluted according to the manufacturer’s instructions.

### Synchronization and breeding

Two to three months after the booster, the heifers were synchronized; they were given 1 mL IM gonadotropin-releasing hormone (GnRH, Gestavet-GnRH, HIPRA) and had an intravaginal progesterone-impregnated controlled internal drug release device (CIDR, Zoetis) inserted. The CIDR devices were removed after five days, and 1 mL prostaglandin was administered to the heifers (D-cloprostenol, Gestavet-Prost HIPRA). A second dose of prostaglandin was administered after 24 hours. Forty-eight hours later, the heifers were given 1 mL IM GnRH and were artificially inseminated using frozen semen from a BVDV-free bull. Heifers were confirmed to be pregnant by transrectal ultrasonography approximately 35 days after artificial insemination and prior to challenge. Only pregnant heifers were used for the BVDV challenges. Non-pregnant animals were excluded from the study.

### Challenge

Twenty-two pregnant heifers were used for the BVDV-1 challenge; fourteen were vaccinated and eight were control animals. Twenty-three pregnant heifers were used for the BVDV-2 challenge; seventeen were vaccinated and six were control animals. Each BVDV challenge was performed at a different period of time. As only pregnant animals could be used to assess transplacental infection, the number of vaccinated and control animals in each challenge was different, depending on pregnancy rates in each group.

At Day 85 of gestation, the pregnant heifers were intranasally challenged with ncp BVDV-1 or ncp BVDV-2 isolates. The challenge viruses had been isolated from aborted fetuses submitted from field cases to HIPRA (Spain and Brazil). The 22 heifers challenged with BVDV-1 strain SKOL each received 10^4^ cell culture infectious dose 50% (CCID_50_) of the virus. The 23 heifers challenged with BVDV-2 strain Iguazú each received 10^5^ CCID_50_ of the virus. The inoculum was administered through a disposable nasal applicator aerosol generator coupled to a syringe. Ten millimeters was administered to each heifer (5 mL/nostril). Animals were appropriately restrained, and the head was kept in an upright position for approximately one minute after inoculation.

### Clinical assessment and sample collection

During the vaccination phase, from the first vaccination until one day before challenge, safety parameters were monitored daily (depression and systemic reactions). During the experimental challenge phase, animals were monitored for clinical signs of BVDV infection (nasal discharge, cough, dyspnea, oral mucosa appearance, depression) from one day prior to challenge and then daily until the end of the study. Rectal temperature was also measured daily from one day prior to challenge to 14 days post-challenge.

Blood samples for BVDV antibody detection by ELISA were obtained before the first vaccination (D0). Blood samples for BVDV serum neutralizing antibody detection (SN) were obtained before the first vaccination (D0), before the second dose (D21), then at 21 days (D42), 2 months (D84), and 5 months after the second dose (D166), as well as on the day of the third dose (D203), 21 days after the third dose (D226), at the challenge (D344), 21 days post-challenge (D365), and just before euthanasia (60-67 days after challenge). Heparinized blood samples were collected to obtain peripheral blood mononuclear cells (PBMCs) to determine BVDV-specific IFN-γ levels before challenge and seven days after challenge. Blood samples for white blood cell (WBC) counts were collected in EDTA tubes one day prior to challenge, on the challenge day, and on Days 3, 5, 6, 7, 8, 10, 14 and 21 post-challenge. Additional samples were collected on Days 4, 9, 11, 12, 13 post-challenge in the BVDV-2 trial. EDTA blood samples were also collected to prepare buffy coats (BC) and investigate the presence of BVDV RNA via RT-qPCR before vaccination, on the challenge day, and on Days 3, 5, 6, 7, 8, 9, 10, 14 and 21 post-challenge. Additional samples were collected on Day 11 post-challenge in the BVDV-2 trial.

Nasal swabs (NS) were collected to detect BVDV RNA via RT-qPCR on the challenge day and on Days 3, 5, 6, 7, 8, 9, 10, 12, 14 and 21 post-challenge.

At approximately 150 days of gestation (60−67 days post-challenge), the heifers were euthanized and their fetuses collected. Each fetus was necropsied. The thymus, brain, liver, Peyer’s patches, and spleen were collected. Fetal tissue samples were tested for the presence of BVDV via a virus titration assay.

### Testing of samples

#### BVDV antibody detection

BVDV antibody levels were determined using a commercial ELISA (HerdCheck BVDV antibody test, IDEXX). A serum neutralization test in MDBK cells in 96-well plates was used to quantitate SN antibodies against BVDV-1 and BVDV-2 using cytopathic virus strains Singer (BVDV-1) and VV-670 (BVDV-2). A constant viral titer (10^2^ CCID_50_) was incubated with two-fold dilutions of sera. Culture plates were incubated for seven days and visually assessed for virus-induced cytopathic effects. Geometric mean titers (GMT) were calculated by way of log_2_ titers.

#### BVDV-specific T-cell immune response

PBMCs were used to quantify BVDV-specific IFN-γ secretion by ELISA in response to stimulation with BVDV-1 or BVDV-2. PBMCs were isolated and counted, the concentration of PBMCs in each sample was titrated to 5 × 10^6^ PBMCs/mL with a mixture of RPMI medium and 10% FBS, and then 100 μL aliquots were transferred to replicates of three wells in a 96-well flat-bottom plate. To each well of one replicate of three wells, 100 μL BVDV-1 recombinant protein (10 μg/mL), BVDV-2 recombinant protein (10 μg/ml), Pokeweed mitogen (10 μg/mL), or a mixture of RPMI medium and 10% FBS (cell culture media) was added. Plates were then incubated at 37 °C in 5% CO_2_ for 96 hours. After incubation, plates were centrifuged at 2500 rpm for ten minutes. From each well, 150 μL of supernatant was transferred to another 96-well flat-bottom plate and stored at −80°C until analyzed. IFN-γ secretion by PBMCs in response to stimulation with BVDV-1 and BVDV-2 was measured via a commercially available ELISA (Bovigam, Ingenasa) in accordance with the manufacturer’s instructions. For each sample, IFN-γ was determined as an IRPC value using Pokeweed mitogen with RPMI medium as a positive and negative control, respectively.

#### White blood cell counts

WBC counts were analyzed using a semi-automated electronic cell counting device (XN-1000 Sysmex, Laboratorio Echevarne, Barcelona, Spain).

#### BVDV quantification by RT-qPCR

EDTA blood samples from heifers were used to prepare BC. Red blood cells were lysed with NH_4_Cl for ten minutes, and a pellet containing the WBC was obtained after centrifuging (2000 rpm, five min.). BC cells were washed and aliquoted to a final volume of 1.5 mL in MEMG. NS collected from heifers post-challenge were placed in 3 mL of virus transport medium. Upon arrival at the laboratory, this medium was stored frozen at −70 °C until the analysis for BVDV RNA.

Buffy coat cells and nasal swabs from heifers were assayed for the presence of BVDV RNA by RT-qPCR with SYBR Green methodology, using primer pairs (PEST-3) 5’-GTG GAC GAG GGC ATG CCC A-3’ and (PEST-D) 5’-TCA ACT CCA TGT GCC ATG TA-3’. Total RNA was extracted from BC and NS using Biosprint 96 One-for-all vet Kit (Qiagen) according to the manufacturer’s instructions. RT-PCR cycle conditions were as follows: 50 °C for 30 min., 95 °C for 15 min., and 40 cycles at 94 °C for 15 sec., 58 °C for 30 sec and 72 °C for 30 sec., and finally 95 °C for 15 sec., 60 °C for 1 min. and 95 °C for 30 sec. Thermocycling was performed using a LightCycler480 (Roche).

#### BVDV titration assay

A virus titration assay was used to quantify BVDV in fetal tissue samples obtained during the necropsy. Fetal tissues were homogenized, diluted to 1/2 (brain, liver, Peyer’s patches, and spleen) or 1/5 (thymus) in MEMG supplemented with streptomycin (250 µg/mL) and ampicillin (125 µg/mL), and inoculated in duplicate into 96-well plates that were seeded with MDBK cells and incubated for four days (37 °C, 5% CO_2_). After incubation, an immunoperoxidase monolayer assay (IPMA) was performed. A BVDV-1 ncp isolate was used as a positive control, and a blank cell culture medium served as the negative control.

### Statistical analysis

Statistical analyses and plots were generated using R (version 4.0.5) and Microsoft□ Excel 2010 (Microsoft corp.). Analyses for the two BVDV challenge trials (BVDV-1 and BVDV-2) were performed separately. Unless otherwise specified, all plots depict the sample mean and standard error. When required, data were log_10_-transformed (i.e., log-WBC) to satisfy assumptions of normality.

Statistical analyses of seroneutralizing antibody levels before and after challenge were performed separately. For both periods, a linear mixed-effects model using the lme implementation in the nlme R package was used. Before challenge, time was included in the model as fixed effect, and the experimental subject was considered a random factor. After challenge, time, group and their interaction were included in the model as fixed effects, and the experimental subject was considered a random factor. The corresponding random intercept models were fitted to the data using restricted maximum likelihood. Correlation between longitudinal observations as well as heteroskedasticity were included in the models when required with appropriate variance-covariance structures. Assumptions were tested graphically (using quantile-quantile and residual plots) for both modeling approaches, and model selection was based on likelihood ratio tests or *a priori* assumptions. For pairwise comparisons, the corresponding estimated marginal means were calculated and compared using the emmeans R package.

The geometric mean of antibody titers was calculated from the endpoint log_2_ titers of the animals in each group. Fisher’s exact test was used to analyze differences in the percentage of animals with neutralizing antibodies between the control and vaccinated groups.

Rectal temperature evolution was analyzed analogously to seroneutralizing antibody levels. The only difference was that time, group and their interaction were included in the model as fixed effects, and the experimental subject was considered a random factor.

Average days with hyperthermia (> 39.5 °C; 23) and the clinical sign score were compared using a Mann-Whitney U test between groups. Differences in WBC count between groups were assessed using a t-test or Mann-Whitney U test according to data normality at each time-point evaluated. Additionally, differences in WBC count within groups (control or vaccinated) were analyzed by ANOVA or Kruskal-Wallis tests according to data distribution. BVDV-specific IFN-γ levels were also compared between challenge day and seven days post-challenge using a Mann-Whitney U test.

The percentage of animals with nasal shedding or viremia per group was analyzed using a Fisher’s exact test. The number of days where BVDV was detected on nasal samples or buffy coats per group was compared using a Mann-Whitney U test.

Finally, a Fisher’s exact test was also used to compare the percentage of positive samples from fetuses between groups. Viral titers obtained from the different fetal tissues were compared between groups using a Mann-Whitney U test.

A significance level *p* < 0.05 was used for all variables evaluated in this trial.

## RESULTS

### Serology

In the BVDV-1 trial, vaccinated heifers showed a significant increase (*p* < 0.05) in neutralizing antibodies at Day 21 post-vaccination. A peak mean antibody log_2_ titer of 8.8 (GMT, 438.5) was reached after the third dose of vaccine on Day 226 (Figure 1). Antibody titers decreased slightly prior to challenge, with a mean antibody log_2_ titer of 8.1 (GMT, 280.4) on challenge day. Control heifers had no neutralizing antibodies prior to challenge. Consequently, vaccinated animals had significantly (*p* < 0.05) higher antibody titers and a higher proportion of seropositive animals compared to the control group from Day 42 to challenge day (D344). All animals had SN antibodies 21 days after challenge, indicating that the challenge was correctly conducted. Antibody titers 21 days post-challenge were also significantly (*p* < 0.05) higher in the vaccinated group compared to the control group.

**Figure 1.**
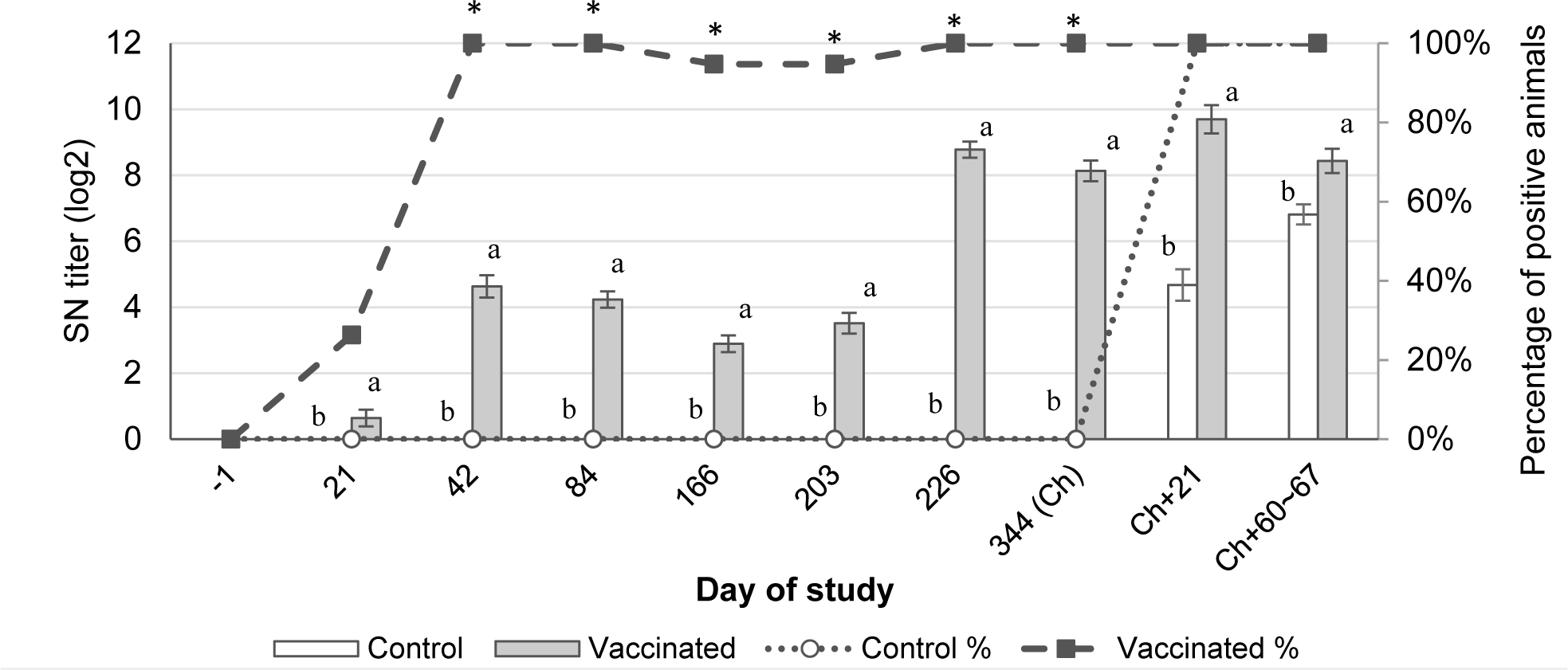
Neutralizing antibodies against BVDV-1 (mean ± SEM) measured by serum neutralization test (bars) plus percentage of positive animals per group (lines) from Day −1 to 407 of the study. Ch. indicates the challenge day. ^a,b^ indicates statistically significant differences between bars (*p* < 0.05).* indicates statistically significant differences between lines (*p* < 0.05).

Similarly, in the BVDV-2 trial, vaccinated heifers also showed a significant (*p* < 0.05) increase in neutralizing antibodies from Day 42 post-vaccination (Figure 2). The mean antibody log_2_ titer at challenge day was 7.5 (GMT, 177.4). Control heifers had no neutralizing antibodies prior to challenge, and all of them had seroconverted 21 days after challenge, indicating that the challenge was correctly conducted. Vaccinated animals had significantly (*p* < 0.05) higher antibody titers and a higher proportion of seropositive animals compared to the control group from Day 42 to challenge day (D344). Antibody titers 21 days post-challenge were also significantly (*p* < 0.05) higher in the vaccinated group compared to the control group.

**Figure 2.**
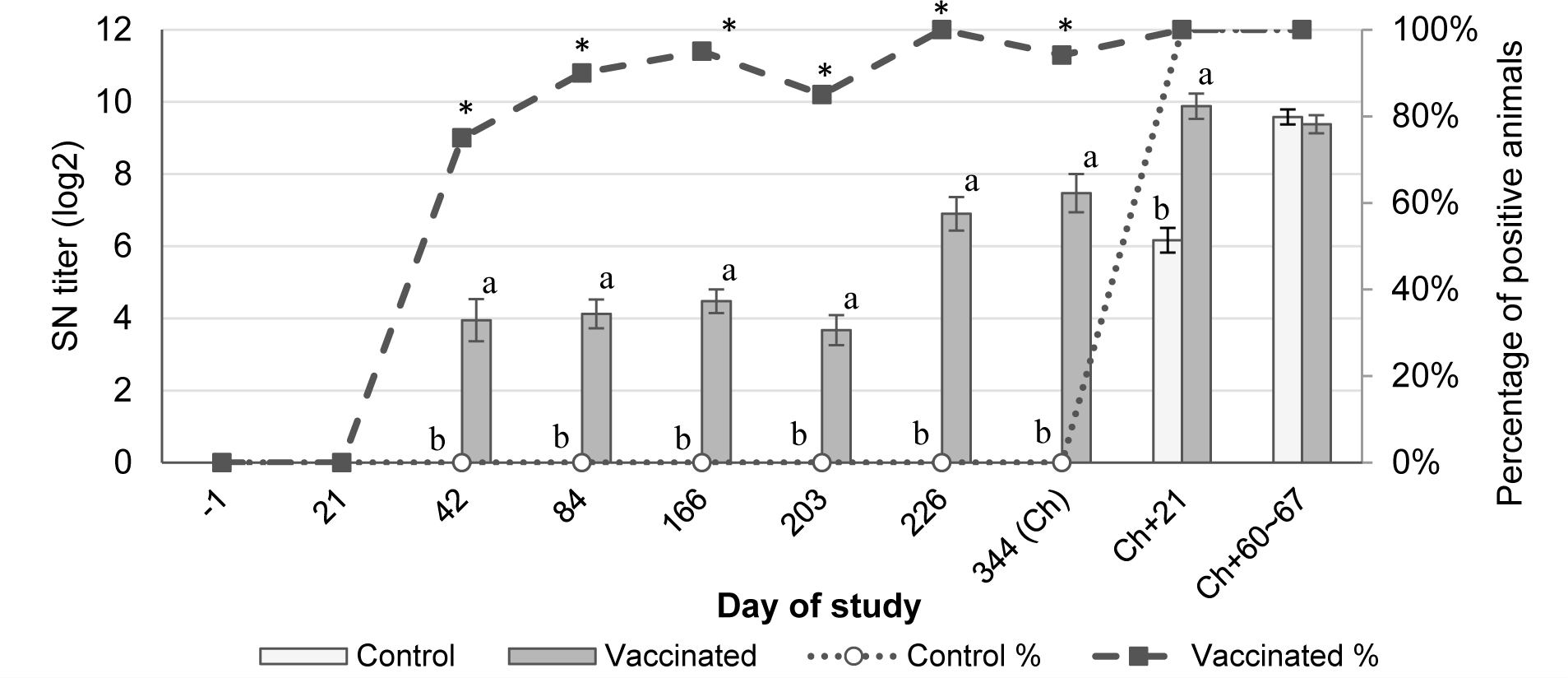
Neutralizing antibodies against BVDV-2 (mean ± SEM) measured by serum neutralization test (bars) plus percentage of positive animals per group (lines) from Day −1 to 407 of the study. Ch. indicates the challenge day. ^a,b^ indicates statistically significant differences between bars (*p* < 0.05).* indicates statistically significant differences between lines (*p* < 0.05).

### Clinical signs and rectal temperature

No relevant adverse systemic effects or injection site reactions were observed during the vaccination phase.

After BVDV-1 challenge, clinical signs in vaccinated and control animals were mild. However, two peaks of rectal temperature were observed in control animals. The first peak was on Day 4, where the average rectal temperature in the control group was significantly (*p* < 0.05) higher than in the vaccinated group. Between Days 5 and 7, rectal temperatures were similar between groups. After that, a second peak of rectal temperature was observed between Days 8 and 11. The average rectal temperature in the control group was significantly (*p* < 0.05) higher than in the vaccinated group at Days 9 and 10 post-challenge (Figure 3). As regards the number of days with hyperthermia (over 39.5 °C; 23), the control group had an average of 1.1 days, which was significantly (*p* < 0.05) higher than the average observed in the vaccinated group (0.2 days).

**Figure 3.**
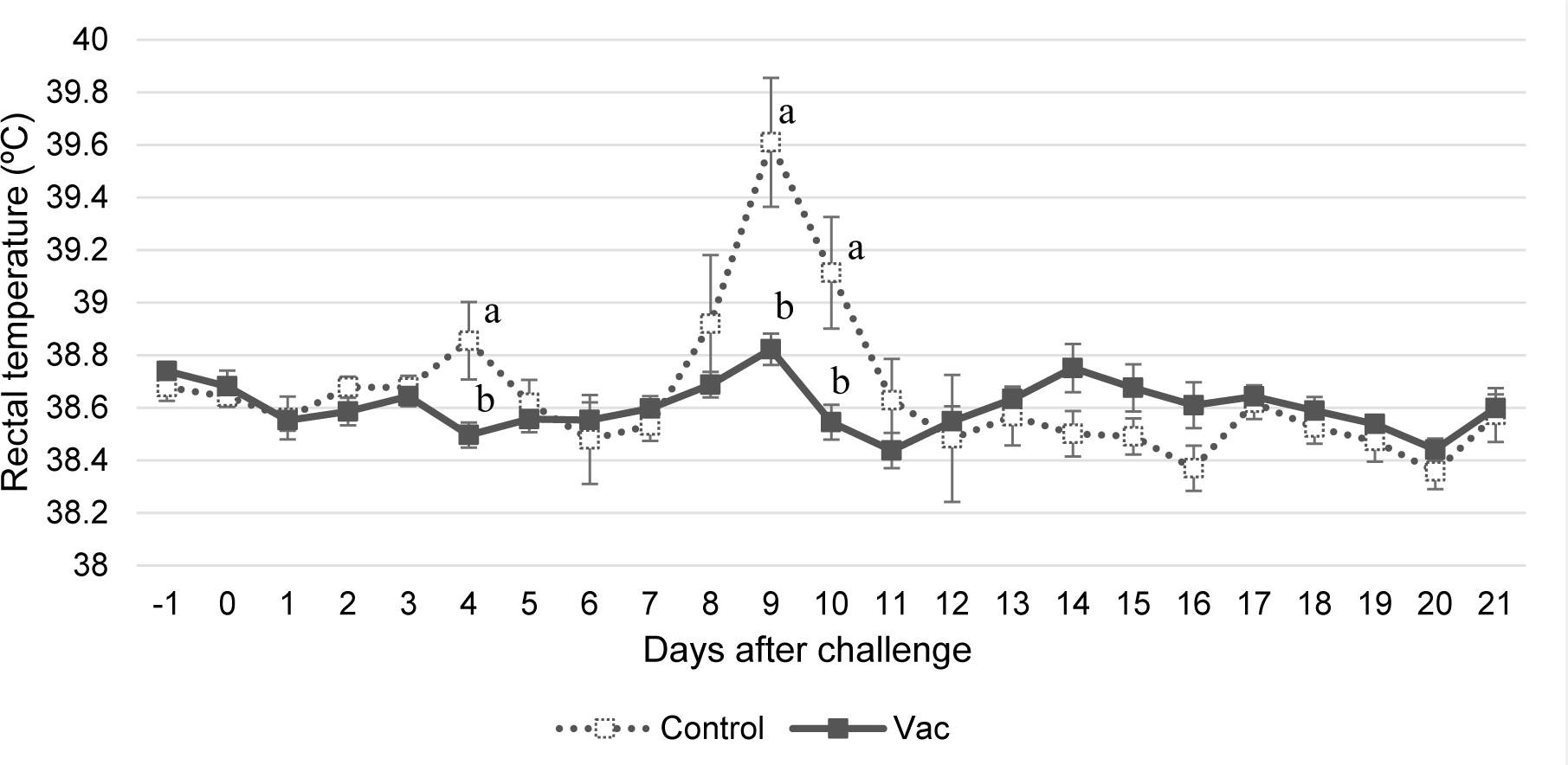
Daily rectal temperatures (Mean ± SEM) per group from one day before challenge until 21 days after challenge with BVDV-1. ^a,b^ indicates statistically significant differences (*p* < 0.05).

In the BVDV-2 challenge trial, clinical signs in vaccinated animals were significantly (*p* < 0.05) lower on Days 9 and 10 post-challenge in comparison with control animals. Mean clinical sign scores in control animals were 1.67 and 1.50 on Days 9 and 10, respectively. In contrast, mean clinical sign scores in vaccinated animals were 0.41 and 0.59 on Days 9 and 10, respectively. The average rectal temperature in the control group was significantly (*p* < 0.05) higher compared to the vaccinated group on Days 4, 5, and 8 post-challenge (Figure 4).

**Figure 4.**
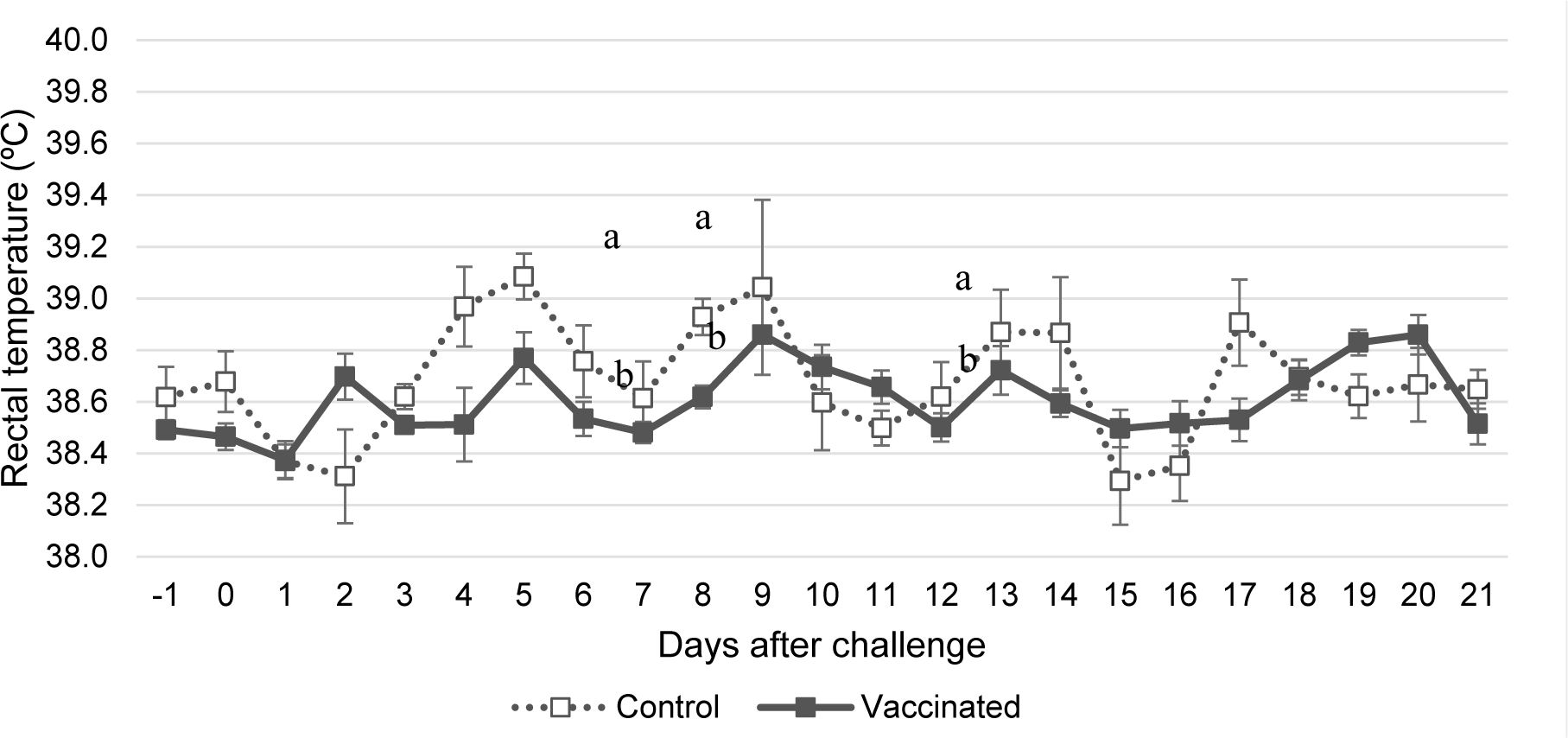
Daily rectal temperatures (Mean ± SEM) per group from one day before challenge until 21 days after challenge with BVDV-2. ^a,b^ indicates statistically significant differences (*p* < 0.05).

### WBC count

After the BVDV-1 challenge, control animals showed a clear and significant (*p* < 0.05) decrease in white blood cells on Days 5 to 7 versus baseline (Days −1 and 0). In contrast, vaccinated animals did not show a significant decrease in white blood cells. The average WBC count in vaccinated animals was significantly (*p* < 0.05) higher than in control animals on Days 5 to 8, 14, and 21 post-challenge (Figure 5). After the BVDV-2 challenge, control animals also showed a significant (*p* < 0.05) decrease in white blood cells on Day 5 post-challenge. In contrast, vaccinated animals again did not show a decrease versus baseline. The average WBC count in vaccinated animals was significantly (*p* < 0.05) higher than in control animals on Days 5 and 6 post-challenge (Figure 6).

**Figure 5.**
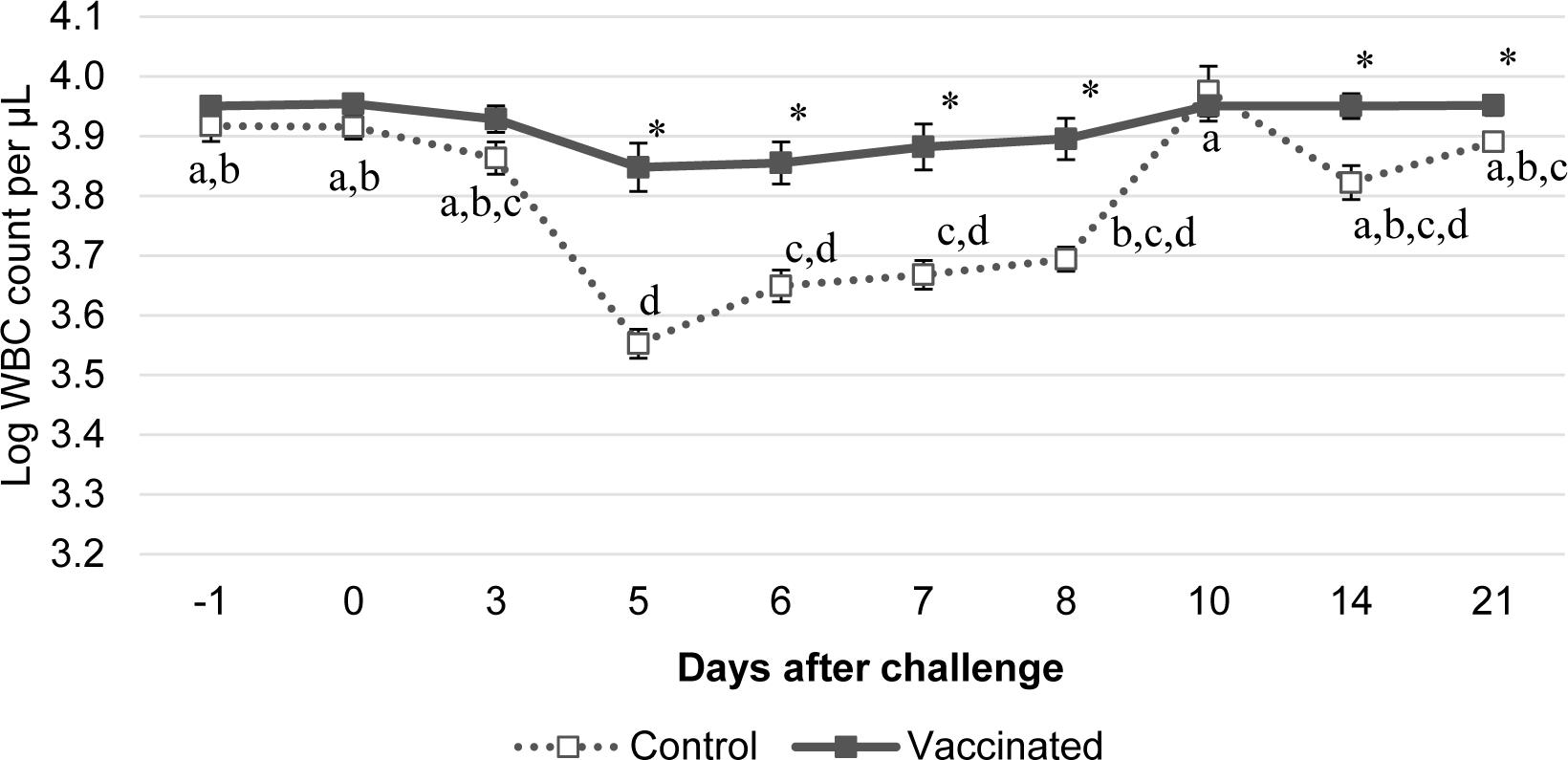
WBC count per group from one day before challenge to 21 days after BVDV-1 challenge. a,b,c,d indicates statistically significant differences within each group when compared with Days −1 and 0 (*p* < 0.05). * indicates statistically significant differences between groups (*p* < 0.05).

**Figure 6.**
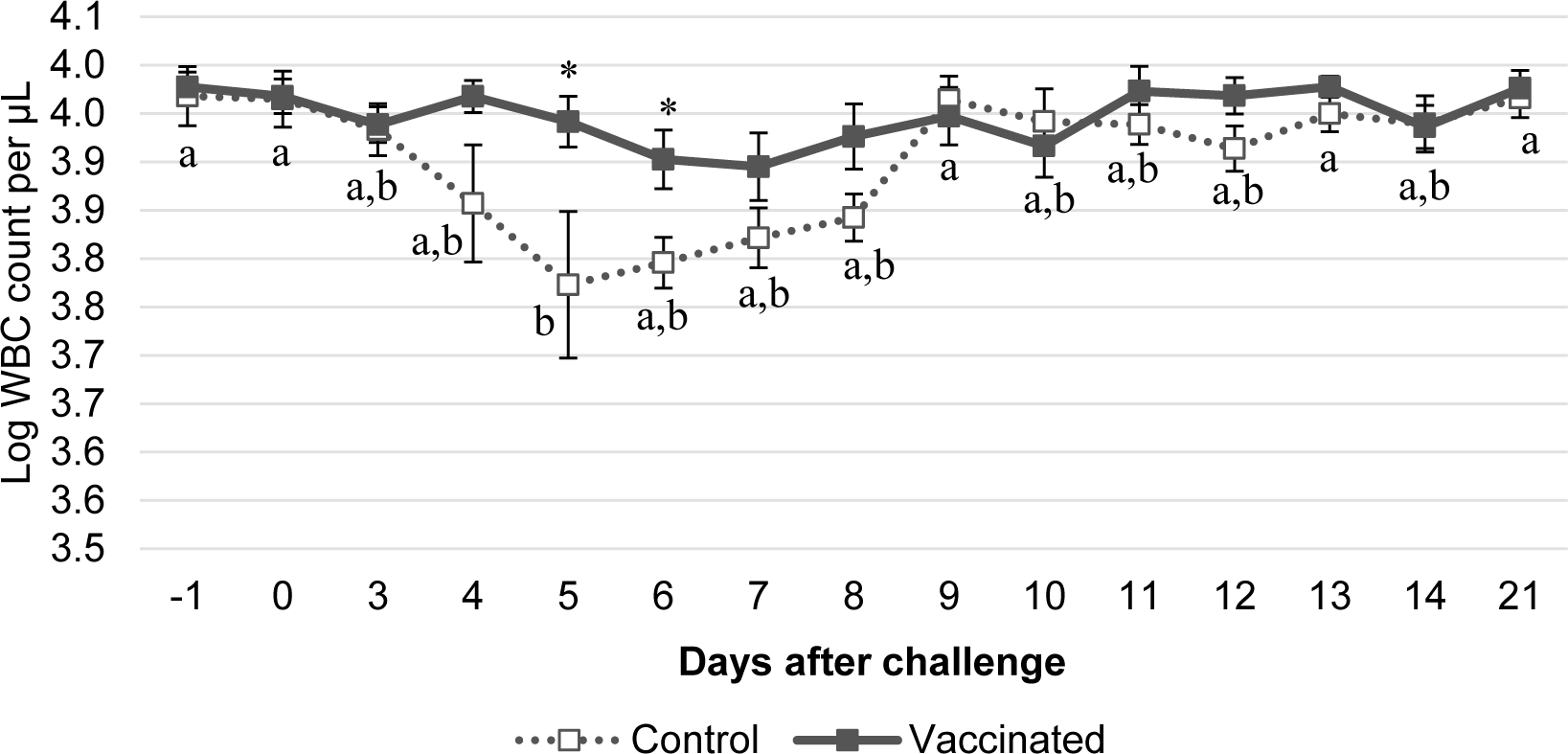
WBC count per group from one day before challenge to 21 days after BVDV-2 challenge. ^a,b^ indicates statistically significant differences within each group when compared with Day −1 and 0 (*p* < 0.05). * indicates statistically significant differences between groups (*p* < 0.05).

### BVDV-specific T-cell immune response

BVDV-specific IFN-γ levels were determined before and after challenge in both trials (BVDV-1 and BVDV-2). On the day of challenge, vaccinated animals elicited significantly (*p* < 0.05) higher BVDV-specific IFN-γ-secreting PBMCs compared to the control group in the BVDV-1 trial (Table 1). This marked difference was also observed seven days later, when vaccinated animals had higher average BVDV-specific IFN-γ-secreting PBMCs compared to challenge day in both trials, while no control animals had IFN-γ expression.

**Table 1.** BVDV-specific IFN-γ levels per group (mean ± SEM) before challenge and seven days after challenge in both challenge trials as detected by ELISA.^a,b^ indicates statistically significant differences between groups within each challenge trial (*p* < 0.05).

| Challenge trial | Treatment group | BVDV-specific IFN-γ levels (IRPC) | |
| --- | --- | --- | --- |
|  |  | Challenge day (Ch) | Ch+7 |
| BVDV-1 | Control | 6.8±3.5<sup>b</sup> | 0±0<sup>b</sup> |
| | Vaccinated | 53.3±15.0<sup>a</sup> | 84.5±28.4<sup>a</sup> |
| BVDV-2 | Control | 0±0 | 0±0<sup>b</sup> |
| | Vaccinated | 18.4±10.8 | 35.3±17.1<sup>a</sup> |

### Nasal shedding

Nasal samples before challenge were found to be negative using RT-qPCR. Post-challenge, BVDV-1 was not detected by PCR in any samples from the vaccinated group, whereas it was detected on nasal swabs in five out of eight control animals. The percentage of positive samples was significantly (*p* < 0.05) higher in the control group than the vaccinated group on Days 8 and 9 post-challenge (Table 2). In terms of duration, control animals shed BVDV-1 for 1.5 days on average, while vaccinated animals exhibited no shedding at all; this marked difference was statistically significant (*p* < 0.05).

**Table 2.**
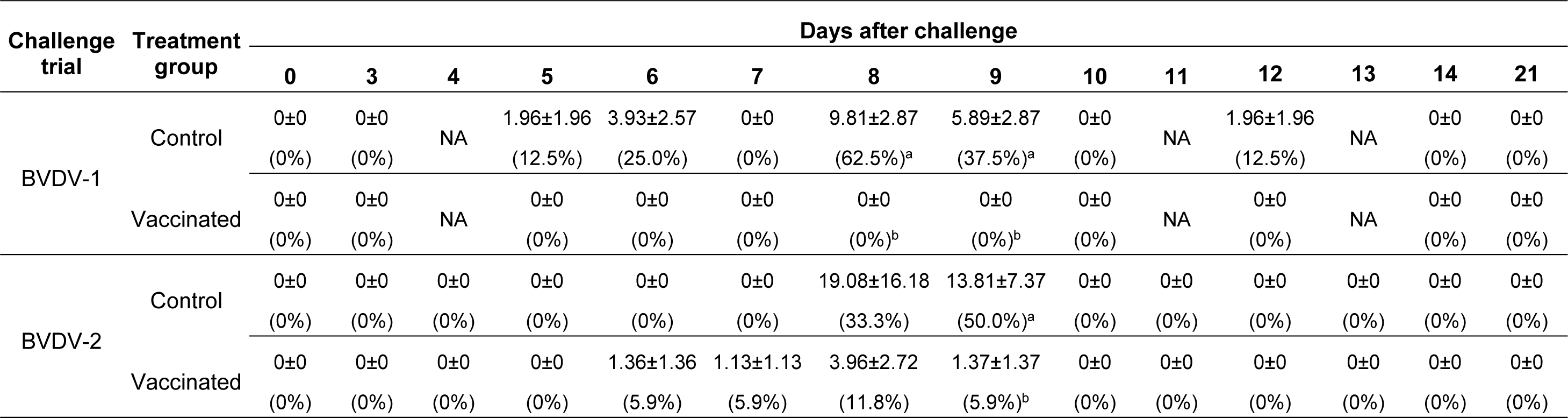
Daily total viral titer per group (mean ± SEM) and percentage of animals shedding BVDV in both challenge trials detected from nasal swabs using RT-qPCR, from challenge to 21 days after challenge. ^a,b^ indicates statistically significant differences (*p* < 0.05).

BVDV-2 was detected by RT-qPCR from Days 6 to 9 post-challenge only in samples from the vaccinated and control groups. Vaccinated animals showed a lower titer than control animals on Day 9 (1.37 vs 13.8 total viral titer, respectively). The percentage of positive samples was significantly (*p* < 0.05) higher in the control group than the vaccinated group on Day 9 post-challenge (Table 2). Over the entire post-challenge period, the percentage of shedding animals was significantly (*p* < 0.05) higher among control animals compared to vaccinated animals.

### Viremia in the heifers

An RT-qPCR assay of BVDV on buffy coat samples was performed to assess viremia in the animals. All control animals (n=8) had viremia after BVDV-1 challenge, while only three out of fourteen vaccinated animals did. These marked differences are shown in Figure 7, where the percentage of viremic animals was significantly (*p* < 0.05) higher in the control group than the vaccinated group on Days 7, 8, and 10 post-challenge. The total viral titer was higher in the control group than in vaccinated animals from Days 7 to 10 post-challenge. In terms of duration, the vaccinated group had a significantly (*p* < 0.05) lower average number of viremic days than the control group (0.2 vs 2.6 days).

**Figure 7.**
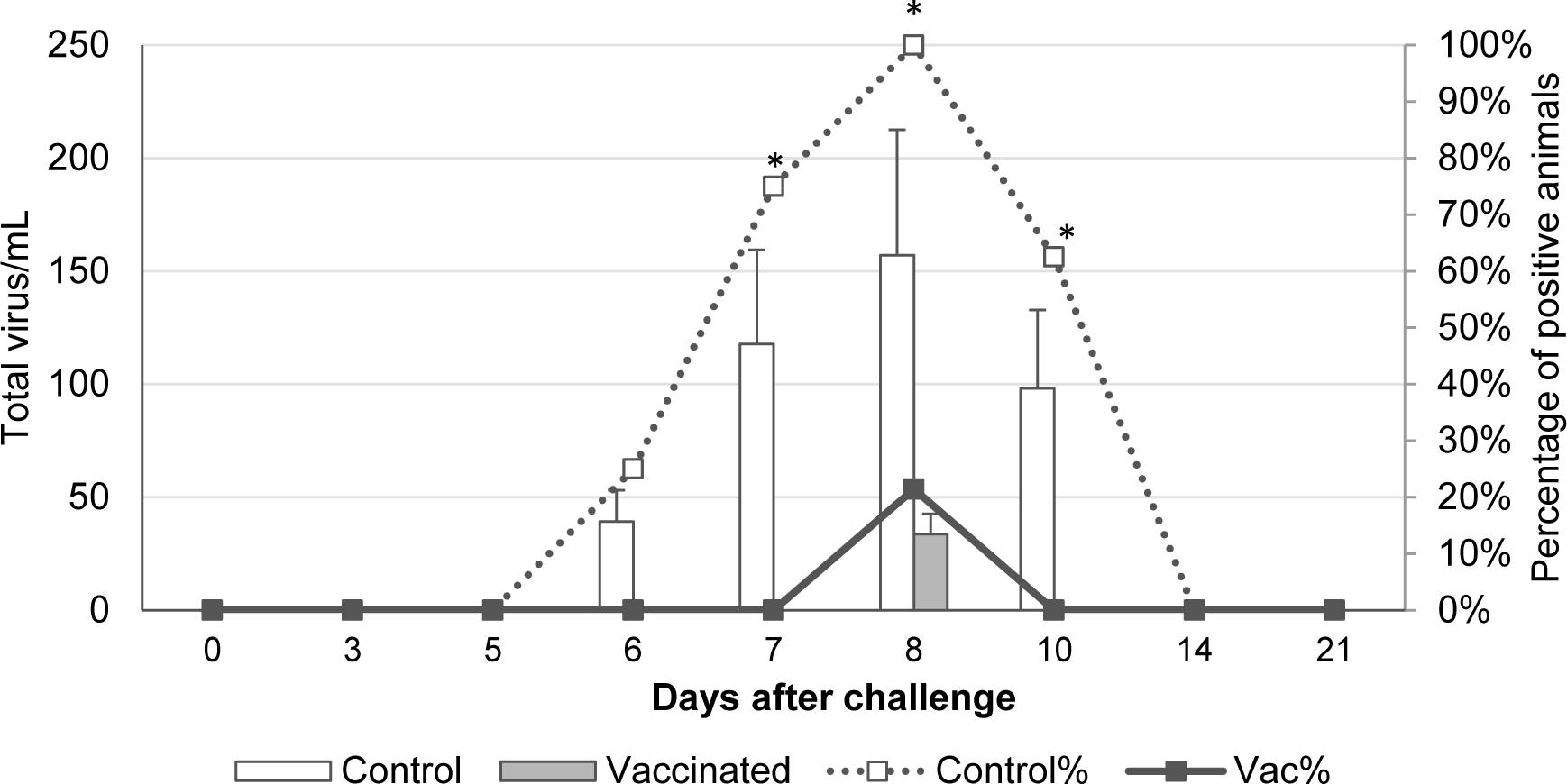
Daily total viral titer per group (mean ± SEM) by RT-qPCR (bars) plus percentage of positive animals per group (lines) from challenge to 21 days after BVDV-1 challenge. * indicates statistically significant differences between lines (*p* < 0.05).

Similarly, all control animals (n=6) had viremia after BVDV-2 challenge, while only three out of seventeen vaccinated animals did. These marked differences are shown in Figure 8, where the percentage of viremic animals was significantly (*p* < 0.05) higher in the control group than the vaccinated group on Days 7 and 8 post-challenge. The total viral titer was higher in the control group than in vaccinated animals from Days 7 to 9. In terms of duration, the vaccinated group had a significantly (*p* < 0.05) lower average number of viremic days than the control group (0.2 vs 2.0 days).

**Figure 8.**
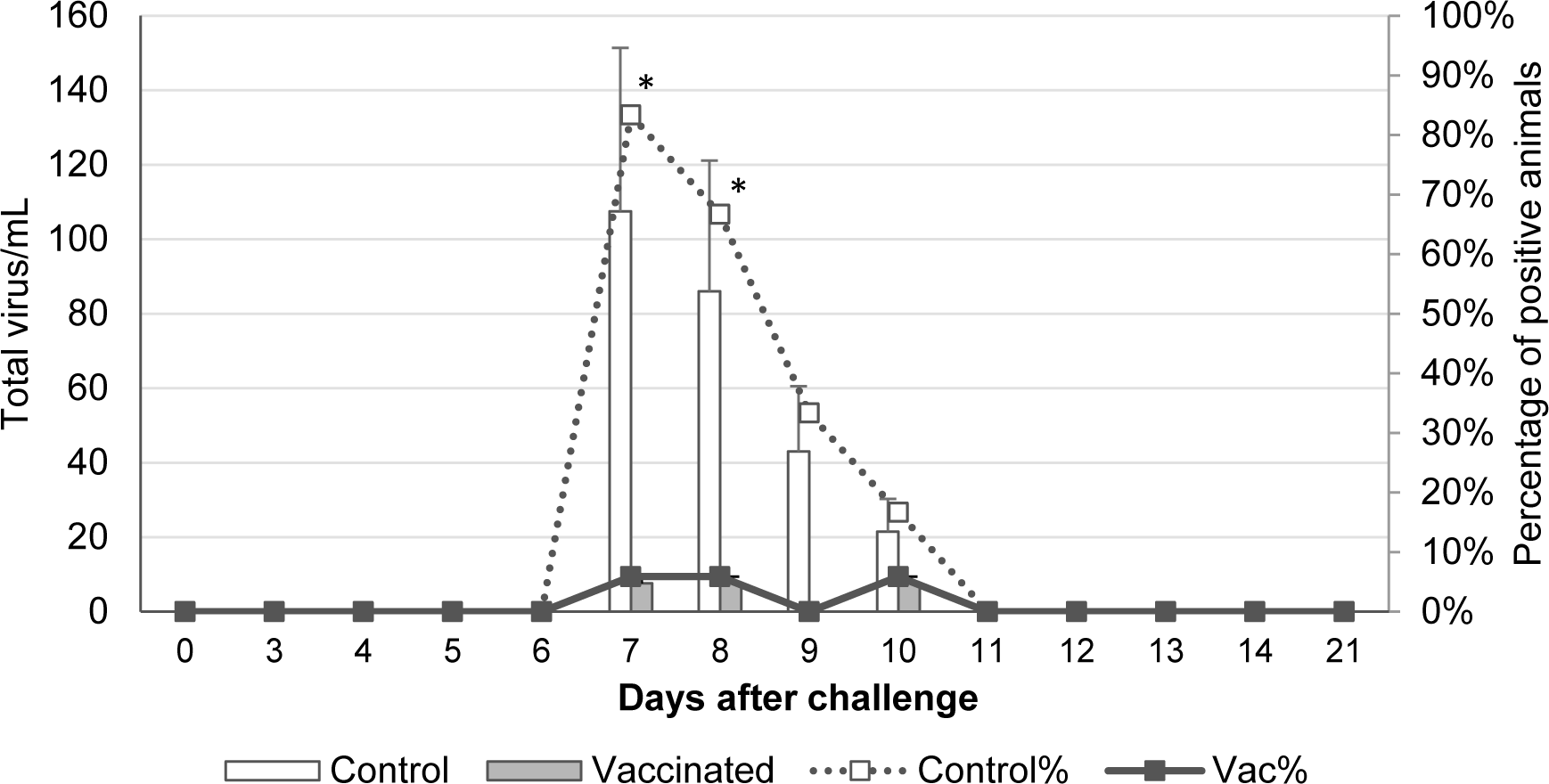
Daily total viral titer per group (mean ± SEM) by RT-qPCR (bars) plus percentage of positive animals per group (lines) from challenge to 21 days after BVDV-2 challenge. * indicates statistically significant differences between lines (*p* < 0.05).

### Transplacental infection

In the BVDV-1 trial, all the control animals had BVDV-1 in the brain and Peyer’s patches taken from fetal samples (n=8). The virus was also detected in the thymus gland and liver in seven of the eight control animals, and in the spleen in six of the eight control animals (Figure 9). In contrast, only one of the fourteen vaccinated animals had BVDV-1 in its fetal samples. Consequently, BVDV-1 was detected in 100% of fetal samples from the control group versus 7.1% in the vaccinated group. These differences were statistically significant (*p* < 0.05). The average BVDV-1 titer in fetal samples was significantly (*p* < 0.05) higher in the control group than the vaccinated group in all tissues collected.

**Figure 9.**
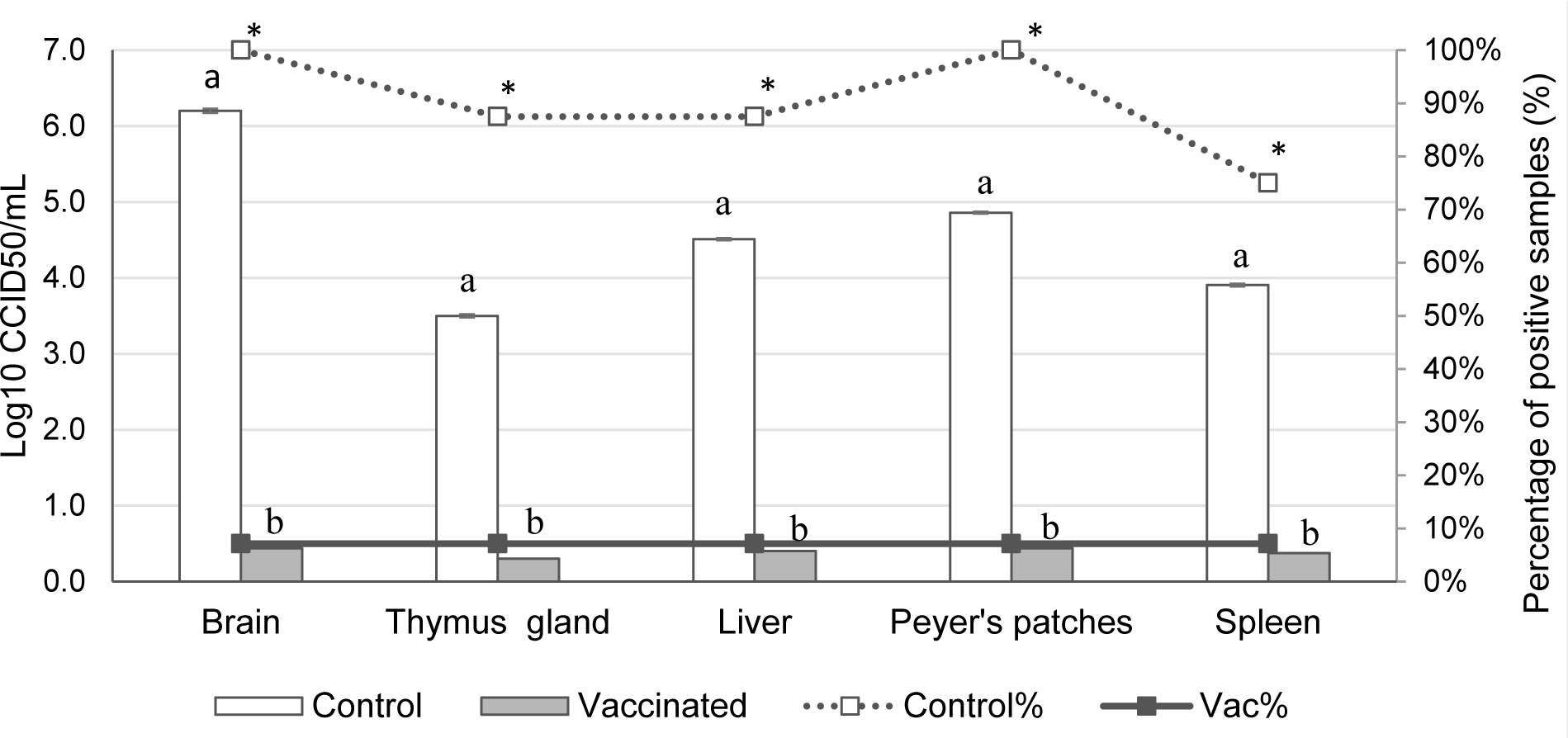
Average BVDV-1 titer (bars) per group of each sample collected from fetuses plus percentage (lines) of positive samples from fetuses. ^a,b^ indicates statistically significant differences between bars (*p* < 0.05). * indicates statistically significant differences between lines (*p* < 0.05).

In the BVDV-2 trial, all the control animals had BVDV-2 in all fetal tissues assessed (brain, spleen, thymus, Peyer’s patches, and liver). In contrast, only one of the seventeen vaccinated animals had BVDV-2 in all tissues. Consequently, BVDV-2 was detected in 100% of fetal samples from the control group versus 5.9% in the vaccinated group. These differences were statistically significant (*p* < 0.05). The average BVDV-2 titer was significantly (*p* < 0.05) higher in the control group than the vaccinated group in all samples collected (Figure 10).

**Figure 10.**
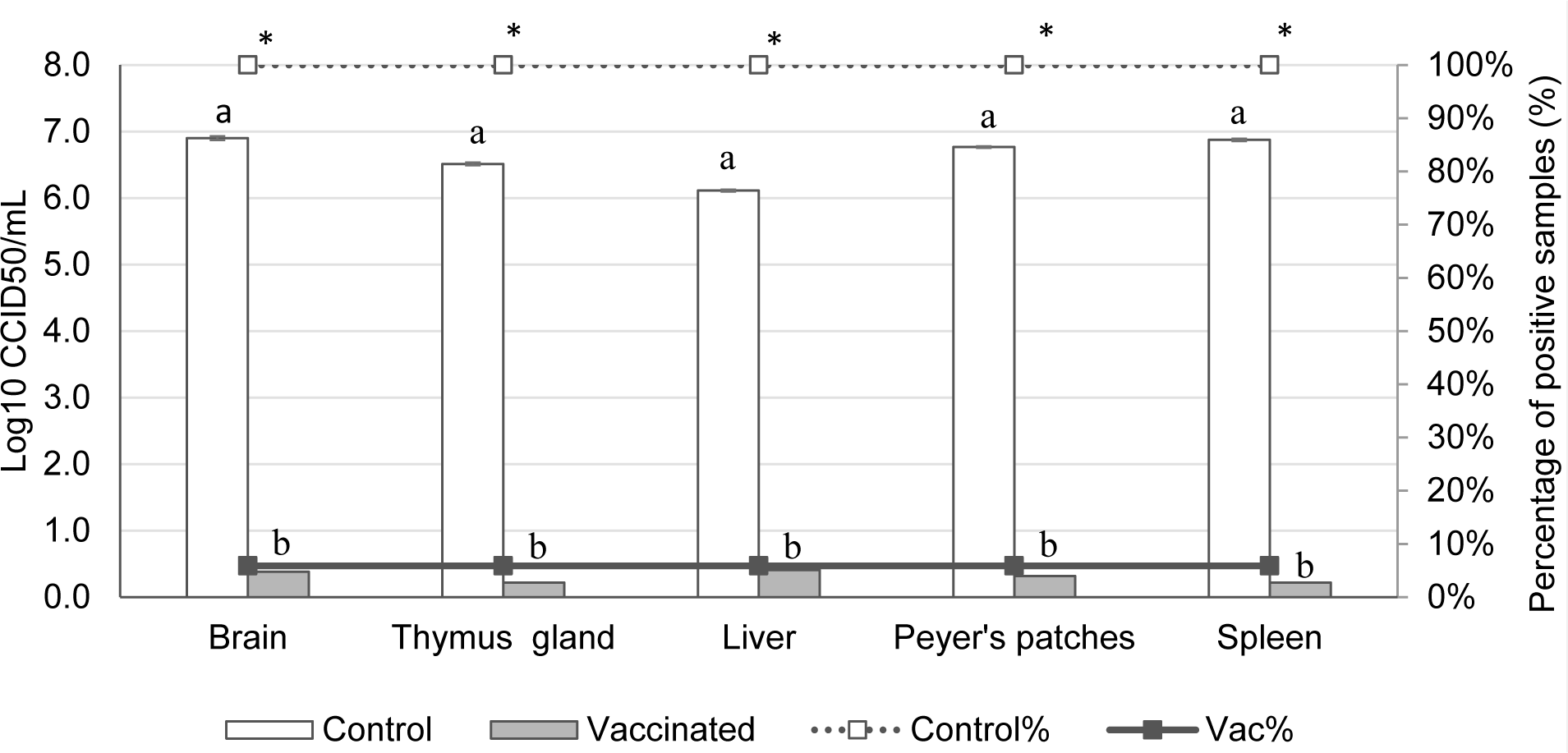
Average BVDV-2 titer (bars) per group of each sample collected from fetuses plus percentage (lines) of positive samples from fetuses. ^a,b^ indicates statistically significant differences between bars (*p* < 0.05). * indicates statistically significant differences between lines (*p* < 0.05).

Overall, DIVENCE administered prior to breeding resulted in 92.9% protection of fetuses from transplacental infection with BVDV-1 and 94.1% for BVDV-2. This results in an overall fetal protection rate against BVDV of 93.5% across both studies.

## DISCUSSION

BVDV vaccination is a valuable tool for disease control and is a common approach adopted in eradication programs, especially in countries with high prevalence (3). Examples include Germany, Ireland, and Scotland, which have adopted vaccination as one of the key measures to control and eradicate BVDV.

However, commercially available vaccines containing only BVDV-1 may not confer cross-protection for BVDV-2 infection (24). This limitation is due to the marked genetic variability between type 1 and 2 BVDV, and a new vaccine must therefore include both genotypes as antigens. In parallel, inactivated and modified-live vaccines (MLV) are used to protect cattle from BVDV infection, but there are concerns about their safety and/or efficacy (25,26). It would therefore be beneficial to combine the immunogenicity of live-attenuated vaccines with the safety of inactivated ones by way of genetic engineering techniques such as subunit vaccines. Subunit vaccines also allow infected and vaccinated animals to be differentiated (DIVA or marker vaccines), which constitutes a new and important tool for veterinarians and farmers to be able to monitor the BVDV status of farms.

DIVENCE is a novel BVDV subunit vaccine containing five antigens – live-genetically modified BoHV-1 (gE-/tk-), live-attenuated BRSV, inactivated PI3 virus, BVDV-1 E2 recombinant glycoprotein, and BVDV-2 E2 recombinant glycoprotein – developed to protect cattle against all these pathogens. This vaccine has been designed to protect against the major pathogens of BRD (BoHV-1, BRSV, PI3, and BVDV-1 and 2) during the first months of life, but also and more importantly to protect fetuses from infection with BVDV-1 and 2. BVDV infection during pregnancy leads to different outcomes, including transient infertility, early embryonic death, fetal mummification, abortion, fetal malformations, birth of PI calves, and congenital infection, depending on the time of viral exposure (20,27). The birth of PI animals is the primary source of BVDV infection and spread within and between farms (3).

The aim of this study was to assess the ability of this vaccine to protect fetuses from transplacental infection. Vaccinated and control pregnant heifers were challenged intranasally with either BVDV-1 or BVDV-2 at 85 days of gestation. Tissues from each fetus were collected and tested for the presence of BVDV via a virus isolation technique. Of the eight fetuses from control heifers infected with BVDV-1, BVDV was isolated from the brain and Peyer’s patches in 100% of the samples. Similarly, of the six fetuses from control heifers infected with BVDV-2, the virus was isolated from all tissues analyzed (brain, spleen, Peyer’s patches, liver, and thymus gland). These results suggest that the brain and Peyer’s patches should be considered the tissues of choice for reliable detection of BVDV in fetal samples. Other authors (27) have also described the brain as one of the most reliable tissues to detect BVDV, as all infected animals had BVDV in fetal brains. Other studies (28) showed that the lungs and spleen are also good fetal samples to assess BVDV infection.

In heifers vaccinated with DIVENCE prior to breeding, an overall fetal protection rate against BVDV infection of 93.5% was observed (92.9% against BVDV-1 and 94.1% against BVDV-2). In studies using ordinary inactivated vaccines, it has been demonstrated that the level of fetal protection conferred is inadequate, ranging from 22%– to 73% (29,30,31). Conversely, several commercial MLV vaccines demonstrated higher efficacy against fetal infection against BVDV-1 and BVDV-2, with most studies reporting more than 85% protection, (28,32,33,34,35). However, safety concerns, particularly in pregnant cattle, are a major issue (36,37,38,39,40,41), which limits the use and applicability of MLV vaccines at herd level and confers a suboptimal herd immunity (20,44). The results obtained in the present study thus demonstrate that a subunit vaccine (DIVENCE) is an effective and safer alternative against fetal infections.

BVDV vaccines should be able to prevent transplacental infection (36), but as stated above, inactivated vaccines have failed to demonstrate an adequate level of fetal protection, which ranges from only 22 to 73% (29,30,31). Nevertheless, the main advantage of inactivated vaccines is their higher safety compared to MLV vaccines. The WOAH does not recommend the administration of MLV vaccines to pregnant cattle (or their sucking calves) due to the risk of transplacental infection. Moreover, MLV vaccines that contain cp strains present a risk of producing mucosal disease in PI animals (36).

Despite the higher efficacy of MLV vaccines compared to inactivated vaccines in the prevention of transplacental infection, the trade-offs are costly. Together with mucosal disease, immunosuppression and fetal infection resulting in abortion and congenital defects have been widely reported in the Americas and Europe after the use of MLV vaccines (36,37,38,39). In addition, there is a potential for viral mutations as the virus replicates *in vivo*, which may result in enhanced virulence (40). Even the use of MLV vaccines in non-pregnant cattle may result in impaired fertility; BVDV from the vaccine has been detected in the ovaries up to one month post-vaccination, which is problematic as it can cause ovarian dysfunction and consequently reduced fertility (37). Similarly, the use of MLV vaccines in peripubertal bulls has also been identified as a risk factor for venereal transmission of BVDV and subfertility. Vaccination with MLV strains of BVDV can result in prolonged viral replication in the testicular tissue of bulls for up to 134 days (41).

As a result, the use of MLV vaccines is often restricted to non-pregnant cattle, even excluding breeding bulls (42), which impacts the protocols that can be applied at the herd level (i.e., transition cows during the voluntary waiting period shortly after calving and before insemination). Furthermore, ordinary multivalent MLV vaccines (including both BVDV and BoHV-1, i.e., non-marker vaccines) have been reported to be abortifacient (43). Accordingly, protocols including this type of vaccine are accommodated and used, as described above. Vaccination in transition cows has also proven less effective in immunizing animals; metabolic disorders are common, resulting in an interaction between impaired nutrient metabolism, oxidative stress, and dysfunctional inflammatory responses, which harms productivity and health (20). One study (44) demonstrated that less than 50% of cows vaccinated during the transition period had increased antibody levels. The strategy of vaccinating cows during the voluntary waiting period may reduce the immune response expected after vaccination.

Conversely, the use of safer vaccines allows different vaccination strategies, such as mass vaccination; the whole herd can be vaccinated at the same time, independently of pregnancy status. The combination of a BVDV subunit and a live BoHV-1 genetically modified vaccine is non-abortifacient, allowing pregnant cattle to be vaccinated with no risk of abortion caused by the vaccine strain, as already demonstrated with the same gE-tk-double-deleted BoHV-1 strain (45). DIVENCE can also be used in young animals for BRD prevention, which allows almost the entire herd to be vaccinated concomitantly.

Mass vaccination has been demonstrated to increase the success of vaccination programs by enhancing herd immunity (46). Similarly, a higher number of doses increases the immunity level within a given population; a single effective round of vaccination may result in an immunity level of 70%, whereas a second round of vaccination can achieve 90% of immunized individuals (46). From a practical perspective, vaccination programs must include a high proportion of the population, but must also uniformly cover the whole herd, as pockets of susceptible individuals can allow a pathogen to persist or recirculate (46). The results presented here mimic the efficacy of fetal protection expected at the individual level if a herd vaccination protocol is used from an early age to prevent BRD as well as thereafter to prevent reproductive disease in heifers and cows. By reducing viral circulation at the herd level through herd immunity, results in terms of fetal protection are expected to be even higher.

To date, only inactivated and MLV vaccines are commercially available; DIVENCE constitutes a third group of BVDV vaccines using different technology, as it contains only recombinant E2 proteins. The structural envelope glycoprotein, E2, is the major immunogenic determinant of the BVDV virion (47). Neutralizing antibodies (NAbs) induced in infected animals are mainly directed against E2 (48). Moreover, E2-specific monoclonal antibodies can neutralize both BVDV-1 and BVDV-2 (49). These findings support the design of an E2-based BVDV vaccine which can distinguish infected from vaccinated animals (DIVA) as a new tool.

The new subunit vaccine (DIVENCE) offers efficacy without the trade-offs of the MLV vaccines, and thus ensuring higher safety. As stated above, the use of these E2 recombinant proteins also implies a new characteristic for BVDV vaccines, as infected and vaccinated animals can be differentiated (DIVA or marker vaccines). Since the vaccine contains only the immunogenic glycoprotein E2, present in BVDV-1 and BVDV-2, vaccination does not induce the production of antibodies against any other protein of BVDV. Therefore, any available commercial kit to measure antibodies against the absent proteins can be used to differentiate infected from vaccinated animals, as was already described with other pathogens such as Aujeszky’s disease virus (22) and BoHV-1 some decades ago.

Glycoprotein E2 is a structural protein of BVDV and the major target for neutralizing antibodies; it thus confers protection after vaccination or infection (50). Conversely, p80 (or NS3) is an immunogenic non-structural protein, often used in commercially available ELISA kits to detect virus exposure (50). Since non-structural proteins are produced while the virus replicates, natural infection or vaccination with MLV vaccines induce the production of p80 antibodies when animals are exposed to the virus. Following this rationale, several attempts have been made to utilize p80 antibodies in combination with live-attenuated vaccines with the aim of validating this combination for the DIVA approach. Although inactivated vaccines should not induce the production of p80 antibodies in vaccinated animals, since the virus does not replicate within the animal, in practice, it has been demonstrated that the use of inactivated vaccines interferes with ELISA monitoring in blood and milk samples (21,51,52).

Antibody detection is the most cost- and time-effective method to identify herd exposure to BVDV (51). Demonstration of BVDV antibodies provides a reliable insight into the level of exposure to BVDV within a group of animals. Diagnosis at the group or herd level is a key part of the assessment stage of a disease control program (50) and may be used to monitor the evolution of seroprevalence over the longer term. However, the use of ordinary BVDV vaccines (both MLV and inactivated vaccines) interferes with interpretation of serological results. ELISA tests are performed in blood or milk samples (i.e., bulk tank milk) to assess exposure and monitor changes in seroprevalence at the herd or group level. Unfortunately, no commercially available MLV vaccines can differentiate infected from vaccinated animals, making the process extremely difficult and limiting the evaluation of BVDV circulation. Likewise, inactivated vaccines have also failed to demonstrate reliable monitoring, as stated above (21,51,52). This new technology applied to BVDV vaccines (recombinant proteins) makes this third group of vaccines a perfect-fit candidate to allow monitoring of BVDV circulation.

The level of neutralizing antibodies induced by vaccination is a good measure to assess vaccine efficacy (53). This study has demonstrated that the administration of DIVENCE induced an increase in neutralizing antibodies against BVDV-1 and 2. In both trials, the mean antibody log_2_ titer was over 7 (1:128) before challenge. Previous studies (54) reported that neutralizing antibodies log_10_ titers higher or equal to 2 (1:100) are important for protection against BVDV infection.

The adaptive immune response to a given pathogen involves a humoral component and a cell-mediated response. Cell-mediated responses are generally characterized by the induction of IL-2, IFN-γ and CD25 labeling (55). A previous study (56) demonstrated that lymphocytes from calves exposed to BVDV in the presence of maternal antibodies produced more IFN-γ after *in vitro* BVDV exposure. These results indicated that calves can develop an antigen-specific T-cell response even in the presence of maternally derived antibodies. In the present study, vaccinated animals had stronger IFN-γ responses to defined BVDV T-cell epitopes when compared to control animals after challenge. Therefore, vaccination with DIVENCE induced a detectable T-cell activation response to BVDV, indicative of a cellular response to BVDV.

Clinical signs after both challenges were mild. Acute BVDV infection has a broad spectrum of clinical presentations, depending on factors such as animal age or the virulence of the BVDV strain. Previous challenge studies also found mild-to-moderate clinical signs of BVDV infection (35). However, despite the mild clinical presentation in this study, control heifers challenged with BVDV-1 and BVDV-2 showed significantly increased mean rectal temperatures post-challenge when compared to vaccinated animals. It can therefore be concluded that DIVENCE reduces the increase in rectal temperature in vaccinated animals challenged with BVDV-1 and/or BVDV-2.

Different BVDV infections, whether subclinical or clinical, can induce immunosuppression, affecting both the innate and the acquired immune system (57). A decrease in total WBC count has been demonstrated in animals challenged with BVDV in other studies (32,58). In the present trial, a clear decrease in white blood cells was observed post-challenge in the control groups (56.8% and 32.7% in the type 1 and 2 BVDV challenge, respectively). In contrast, the WBC count in vaccinated animals did not differ from baseline. The immunosuppressive effect of BVDV is associated with increased severity of co-infections and secondary infections with other pathogens in the field (8,59). DIVENCE thus prevents the immunosuppression of vaccinated animals post-challenge, which helps to protect these animals from other possible infections.

There is evidence of immunosuppression as an adverse effect associated with MLV vaccination due to the strain of BVDV in the vaccine. Similarly, the use of MLV BVDV vaccines has been reported to possibly enhance BRD problems in fattening animals (57). These MLV strains may not differ much in the way they interact with other wild pathogens, or they may be inadequately attenuated (46); BVDV has been shown to correlate positively with BRSV in the case of BRD problems in dairy and beef cattle (9). Other publications show that BVDV worsens the severity of BRSV outbreaks (60) and facilitates or increases its replication (61).

After challenge, the virus was isolated on buffy coat samples from all control heifers challenged with BVDV (type 1 or 2); in contrast, only a few vaccinated animals had viremia, and titers were low. Viremia in pregnant animals allows the virus to cross the placenta and infect the fetus (62). Several researchers (63,64) have found a relationship between the degree of viremia (higher viral titers in blood) and the severity of the disease (a more severe clinical picture). The reduction in the number of days of viremia and in the total viral titer observed in vaccinated animals in this study will help to prevent transplacental infection and birth of PI offspring.

BVDV virus is shed through different excretions and secretions of the infected animals (i.e., nasal and ocular discharge, oral fluids, urine, semen, colostrum/milk, and feces) (62,65). Direct nose-to-nose contact between a PI animal and a susceptible animal has been described as the most plausible route of BVDV transmission (66). Accordingly, nasal secretion is one of the transmission routes of the virus. After BVDV-1 challenge, nasal shedding was prevented in all vaccinated animals, while five of the eight control animals shed virus with a mean total viral titer of 9.81. Conversely, following BVDV-2 challenge, shedding was detected in both vaccinated and control animals. Only three of the seventeen vaccinated animals shed virus, with a maximum mean total viral titer of 1.37. This compares to four of the six control animals which shed virus, with a mean total viral titer of 19.08. The marked difference in the percentage of animals excreting virus between the two groups implies that control animals spread larger amounts of virus into the environment. A reduction in viral shedding decreases horizontal transmission and the rate of new BVDV infection on the farm (67).

The main objective of the study was to demonstrate the efficacy of DIVENCE in protecting against transplacental infection; in addition, a reduction in hyperthermia, leukopenia, nasal shedding, and viremia was consistently observed in vaccinated animals, as compared to the results of the control groups. The reduction in all these parameters is of primordial importance. DIVENCE has therefore been demonstrated to reduce clinical (i.e., hyperthermia) and subclinical (i.e., immunosuppression and viremia) signs following BVDV infection, while also reducing the capacity of the virus to spread within a population in two ways:

1- by conferring fetal protection and thus reducing the likelihood of new PI animals being born (vertical transmission), and
2- by preventing (BVDV-1) or reducing (BVDV-2) nasal shedding and thus limiting horizontal transmission.

To date, ordinarily, the choice of BVDV vaccine has primarily been between live-attenuated vaccines, which offer higher efficacy with a trade-off in safety, and inactivated vaccines, which constitute a safer but less effective approach to BVDV prevention. DIVENCE has been shown to offer both high efficacy in fetal protection as well as high safety, as the biological risks associated with MLV vaccines are not applicable. In addition to this, since the vaccine contains BVDV-1 and BVDV-2 recombinant proteins, this new technology applied to BVDV vaccines adds the extra benefit of allowing infected and vaccinated animals to be differentiated, making DIVENCE a strong candidate for the control of BVDV.

## Acknowledgments

The authors would like to thank Manuel Cañete for the statistical help analyzing the data of this paper, as well as all personnel at HIPRA Scientific, particularly Sara Baila, Laura Feixas and Ricard Martín, for the help performing the study.

